# Statistical framework for assessing heterogeneous sensitivity of viruses to ultraviolet, ozone, and free chlorine

**DOI:** 10.1101/2025.01.20.633964

**Authors:** Miina Yanagihara, Patrick Smeets, Fuminari Miura, Shotaro Torii

## Abstract

Disinfection is key to controlling the infection risk caused by viral contaminations. Recent disinfection guidelines often refer to a single virus resistant to the disinfectant of interest, despite a large variation in sensitivity to disinfectants between viruses or even strains within the same species. Here, we show a statistical framework to integrate multiple experimental datasets and measure the variation in sensitivity to disinfectants across different virus species by a parametric distribution, termed the disinfectant sensitivity distribution. To illustrate this framework, we collected 37, 9, and 28 species-dependent inactivation rate constants for ultraviolet (UV), ozone, and free chlorine from systematic reviews, and constructed the sensitivity distributions for those disinfectants. Our approach incorporated the uncertainty in individual inactivation rate constants using a Bayesian framework. The estimated sensitivity distributions suggested that 93.0% (95% CrI: 84.4–97.6%), 99.4% (95% CrI: 86.7–100%), and 90.3% (95% CrI: 78.2–96.6%) of the examined virus species for UV, ozone, and free chlorine are expected to be inactivated by 4-log if the disinfectant dose complies with the values recommended by USEPA Guidelines for 4-log inactivation, respectively. The proposed approach provides a reasonable extrapolation of observed inactivation kinetics to untested viruses and tools for more transparent risk assessments.

**Graphic for Table of Contents (TOC)/Abstract Art:** 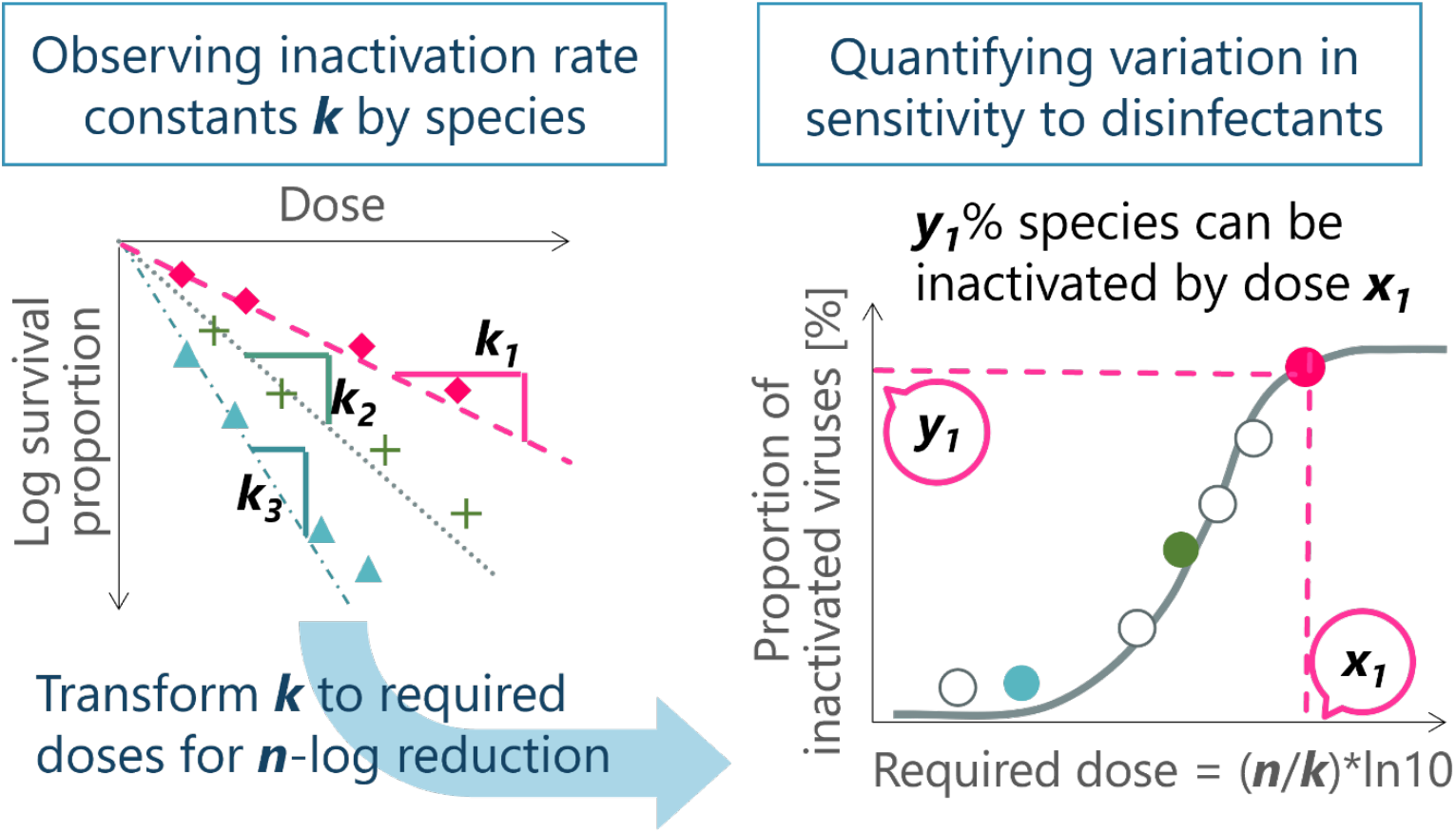

**One sentence summary:** We propose a statistical framework to quantify the variation in sensitivity to common disinfectants across virus species synthesizing disinfection data.

## Introduction

Disinfection is the key component in controlling microbial risks in water, food, air, and other environments. Pathogenic viruses are inactivated with exposure to disinfectants such as free chlorine, ozone, and ultraviolet (UV) irradiation, and those disinfection technologies have been widely applied to various industries e.g., (waste)water treatment processes, healthcare facilities, pharmaceutical engineering, and food industry.^1^ Each disinfectant has a different mode of action,^2^ and thus does not always provide high inactivation efficacy against all viruses, which possess various structures of genome and protein. To compare the disinfection efficacy across different types of viruses, the inactivation rate constant (often denoted as *k*), or its reciprocal, required disinfectant dose (e.g., UV fluence, and CT values) for a given level of inactivation, has been used as a common indicator. This estimate can be obtained through fitting the Chick-Watson model, where dilution coefficient is assumed to be one, to disinfection experiment data.^3,4^

Recent disinfection studies have shown that there is large heterogeneity in sensitivity to a disinfectant across different types of viruses, between species, genotypes, and even strains, under identical experimental conditions.^5–14^ When assessing a virucidal efficacy of a disinfection process in water treatment, most guidelines refer to a single virus that is known to be resistant to disinfectants and/or clinically important, aiming to safely cover viruses with lower sensitivity. For example, the United States Environmental Protection Agency (USEPA) Guidance manuals set hepatitis A virus as a reference for free chlorine (with a safety factor of three).^15^ However, recent studies have suggested the existence of more resistant viruses (e.g. coxsackievirus B5), and the research community is debating whether recent data on coxsackievirus B5 should be incorporated as a worst-case scenario in current guidelines for the treatment of drinking water and wastewater.^16–18^

Despite its practicality, there are two intrinsic limitations in this “single reference” approach. First, it is impossible to test all (nearly infinite number of) newly identified viruses. As depicted in the above example of coxsackievirus B5, new resistant viruses would always arise over time. This is because, as research advances, more experiments on untested viruses will be conducted, leading to sequential updates of the most resistant reference virus. Second, even if we follow the threshold value recommended by the reference viruses’ experimental data (e.g., CT values for 4 log reduction), it is unclear how much proportion of virus species, including untested ones, existing in the environmental sample of interest would be inactivated under such an inactivation condition.

The same motivation – how to extrapolate the observed data on tested organisms to untested ones – has been investigated in the field of ecotoxicology and ecological risk assessment. This approach, called species sensitivity distribution,^19–22^, assesses the ecological risk of chemicals by estimating the proportion of species affected by a particular chemical. Typically, a parametric distribution is fitted to acute or chronic toxicity data of multiple species (e.g., ECx values, the effect concentration at which the (toxic) outcome of interest is observed for x%) to obtain the distribution of sensitivity to the chemical of interest at a community level. Subsequently, the fifth percentile value is typically used to derive benchmarks such as a predicted no effect concentration (PNEC). In the similar manner, we argue that disinfection experiment data can be summarized to extrapolate observations at an individual species level to the assemblage-level disinfectant sensitivity distribution.

In this study, we propose a statistical framework to describe various viruses’ sensitivity as a distribution by synthesizing available datasets on inactivation rate constants. To illustrate this framework, we collected disinfection experiment data on UV, ozone, and free chlorine from existing systematic reviews^3,23,24^ and inferred disinfectant sensitivity distributions for each disinfectant. We also discuss how the inferred sensitivity distributions can be used for translating a required log reduction value into the proportion of potentially inactivated viruses, aiming to better summarize the available evidence on disinfectants, with an example of comparisons with disinfection guidelines.

## Materials and Methods

### Data collection

We collected 220, 31, and 82 reported values of inactivation rate constants for UV (mJ^-1^ cm^2^), ozone and free chlorine (mg^-1^ min^-1^ L), respectively, from the systematic reviews,^3,23,24^ which include screening processes to assess if the disinfectant dose is precisely determined. These datasets were either standardized under specific pH and temperature conditions or collected under similar experimental conditions, as detailed in the **Appendix**, to capture the variation in rate constant due to biological differences.

The collected data were then aggregated by viral species, based on the taxonomy proposed by the International Committee on Taxonomy of Viruses.^25^ We took the mean and standard deviation (SD) of the reported values^26^ for each species when multiple observations were available for the same species, otherwise we used only the mean. The dataset consists of both enteric viruses specified either by USEPA or WHO guidelines^27,28^ and the others (e.g., bacteriophages, and non-human mammalian viruses etc.).

Subsequently, the inactivation rate constants and the guideline values for three disinfectants (i.e., values recommended by the current USEPA Guidance manuals for water disinfection^15^) were transformed into a unit of dose, for a given n-log reduction value (n = 2, 4, or 6) (mJ cm^-1^ for UV, mg min L^-1^ for ozone and free chlorine). The guideline for water disinfection serves as a motivating example to illustrate the relationship between the single reference value and the framework proposed in this study because the analyzed data mainly consist of enteric viruses or surrogate viruses for water treatment processes, according to the systematic reviews.^3,23,24^ A more detailed procedure of this data processing is described in **Appendix**.

### Estimating disinfectant sensitivity distributions

The disinfectant sensitivity distributions for UV, ozone, and free chlorine were estimated by fitting three parametric distributions to collected inactivation rate constants using a likelihood-based approach.^29,30^ We opted for a log-normal distribution in the main analysis, which has been widely used in estimating SSDs for ecological risk assessment,^31,32^ and the Weibull and gamma distributions as alternative candidate distributions.

A Bayesian framework was employed to estimate the parameters of examined distributions (see **Appendix** for details). This method handles the uncertainty in the observed inactivation rate constants as interval data, assuming that the individual observation is uniformly distributed over the interval defined by mean ± 2 SD. As inactivation rate constants and the guideline values were defined for a given n-log reduction, we estimated the disinfectant sensitivity distributions for each combination of the three disinfectants and n-log reduction values.

The goodness of fit of each model was assessed using Widely Applicable Information Criterion (WAIC) and Leave-One-Out Information Criterion (LOOIC). 95% credible intervals were obtained using posterior samples of Markov Chain Monte Carlo (MCMC).

### Proportions of potentially inactivated virus species

The disinfectant sensitivity distributions provide a proportion of potentially inactivated species for a given disinfectant dose and a given target of n-log reduction. We compared the proportion of species expected to be inactivated by disinfectant doses corresponding to the guideline values for UV, ozone, and free chlorine. Additionally, the required doses of the three disinfectants to inactivate certain proportions of virus species (i.e., doses at 95th, 99th, and 99.9th percentiles) were estimated, when 4-log reduction was aimed.

## Results

### Data description

Our processed data contained the reported inactivation rate constants for UV, ozone, and free chlorine, for 37, 9, and 28 species, respectively. The histograms of transformed mean inactivation rate constants are shown in **Figure S1** in **Appendix**. The mean and SD values obtained for each virus species were employed for the subsequent analysis.

### Estimated disinfectant sensitivity distributions and proportion of viruses inactivated

Disinfectant sensitivity distributions were estimated for UV, ozone, and free chlorine using a lognormal distribution (**Figure 1**). **Figure 1** illustrates the relationships between the required dose of each disinfectant to achieve 4-log reduction and the proportion of potentially inactivated species. Additionally, Weibull and gamma distributions were employed for each combination of the target n-log reduction and disinfectants (**Figure S2–S4**). The estimated three parametric distributions for the same disinfectant and n-log reduction mostly overlapped each other, but the estimated uncertainty range around the upper and lower tails of the distributions varied because of their different functional forms. The estimated parameters and goodness-of-fit of each distribution for 4-log reduction are summarized in **Table S1**. The computed WAIC and LOOIC for each disinfectant sensitivity distribution showed small differences, suggesting that all models described the observed data nearly equally well.^33^ Based on this finding, hereafter the following discussion will focus on the results of lognormal sensitivity distributions. The transformed values of disinfectant doses recommended by USEPA Guidance manuals were 186 mJ cm^-2^, 0.532 mg min L^-1^, 3.0 mg min L^-1^, for UV, ozone, and free chlorine, respectively. Our analysis showed that 93.0% (95% credible interval (CrI): 84.4–97.6), 99.4% (95% CrI: 86.7–100), and 90.3% (95% CrI: 78.2– 96.6) of virus species were expected to be inactivated by 4-log by UV, ozone, and free chlorine, respectively. We found that *Enterovirus A* could not be inactivated by free chlorine at this dose level. In addition, **Figure 1** also provides an implication of the coverage of viruses expected to be inactivated when a required dose level is determined referring to a single species. For example, if disinfection is performed at the level of doses that achieves 4 log-reduction of *Emesvirus zinderi*, a species including bacteriophage MS2, only 33**–**78% of species would be inactivated.

**Fig 1.**
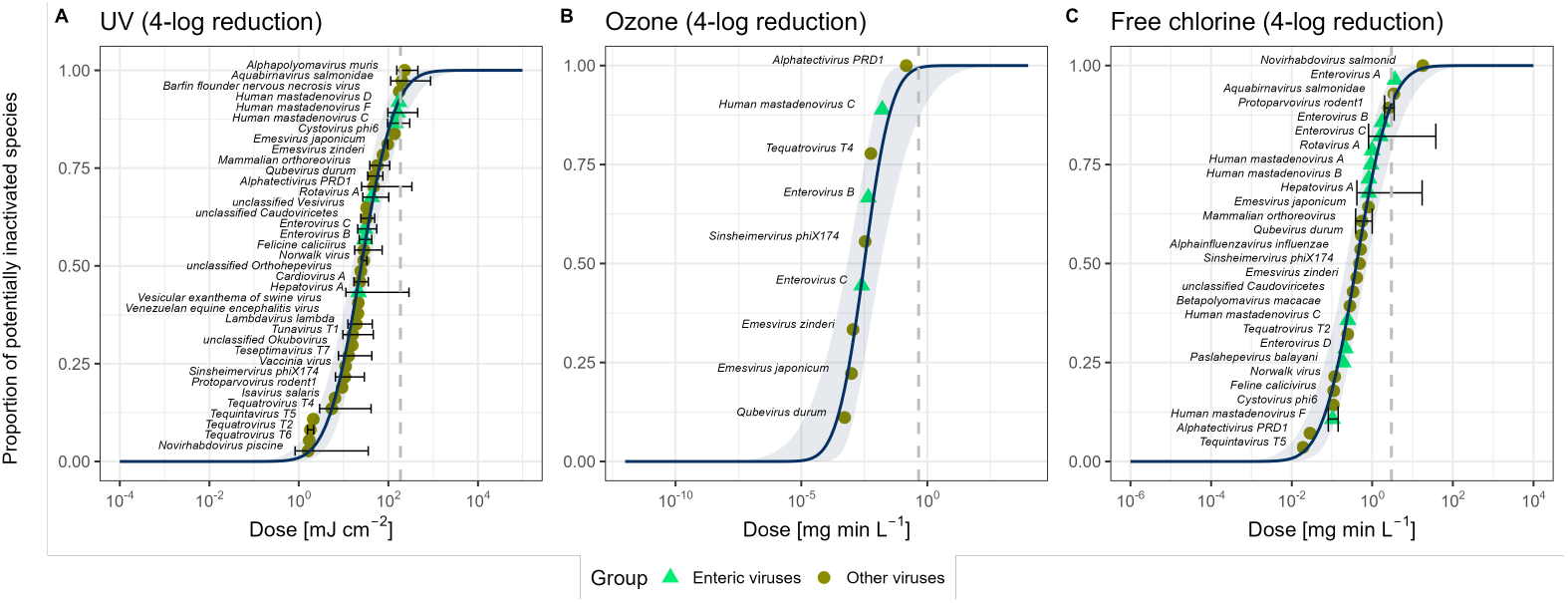
Disinfectant sensitivity distributions of viruses for three disinfectants where 4-log reduction is targeted: UV (A), Ozone (B), and free chlorine (C). Each point and error bar represents the transformed mean inactivation rate constant and the range of two standard deviations (where computable) for each virus. Triangle and green plots represent enteric viruses, and darker circles plots represent other viruses. The species’ names are shown next to each point. The solid lines are the estimated disinfectant sensitivity distributions, and the shaded areas indicate 95% credible intervals computed using posterior samples. The dashed lines represent the recommended dose by USEPA Guidance manuals for each disinfectant.

### Comparison of different guidelines and reference values

In this analysis, disinfectant sensitivity distributions were estimated by changing the target n-log reduction value (n = 2, 4, 6). **Figure 2A** illustrates the estimated UV (log-normal) sensitivity distributions, and shows that the distribution curves shifted in parallel to the higher range of the dose while the curves maintained their shapes (i.e., linear scaling of the x-axis) as the log reduction values increased. See **Figure S2–S4** for all combinations of the disinfectants, log reduction values, and distributions. We also compared the distributions of the proportion of species potentially inactivated by the recommended doses of disinfectants by USEPA guidance when 4-log reduction is aimed (**Figure 2 B–D**).

**Fig 2.**
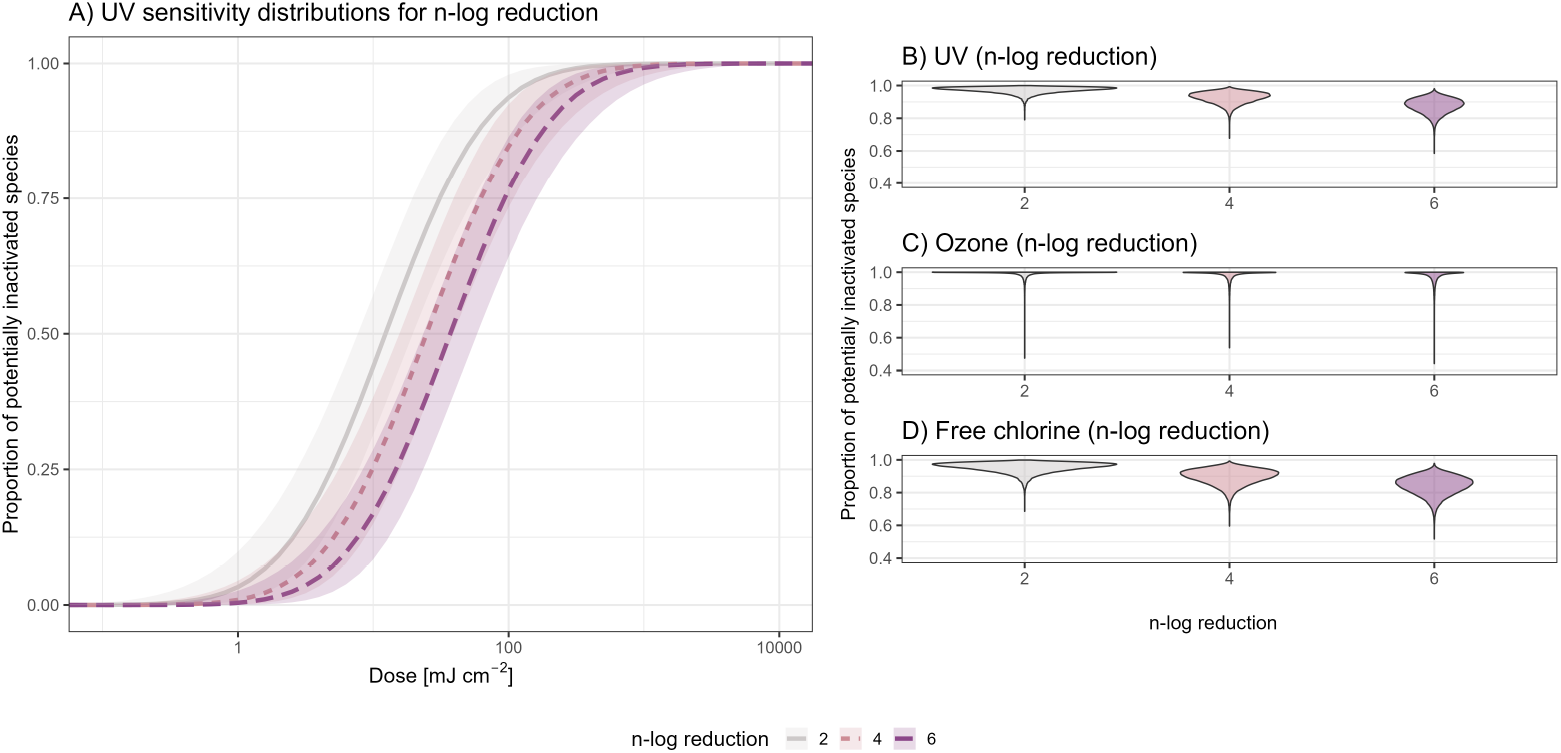
Comparison of the estimated UV sensitivity distributions between different target log reduction values (A). Light gray, light red, and purple plots represent 2-, 4-, 6-log reduction, respectively. Posterior distributions of the proportion of viruses expected to be inactivated by the recommended dose levels by USEPA guidance for UV (B), ozone (C), and free chlorine (D), when 4-log reduction is targeted.

Figure 3. shows the required dose of the three disinfectants to inactivate 95%, 99%, and 99.9% of virus species as posterior distributions (the estimated values are summarized in **Table S2**). For instance, to inactivate 95% of virus species at 4-log reduction, the required dose of disinfectants would be 2.34×10^2^ mJ cm^-2^ (95% CrI: 1.28×10^2^–5.42×10^2^), 0.08 mg min L^-1^ (95% CrI: 0.02–3.02), and 5.12 mg min L^-1^ (95% CrI: 2.37–16.2) for UV, ozone, and free chlorine, respectively. As **Figure 3** depicts, the higher target log reduction value or the proportion of the inactivated species led to an exponential increase in the required disinfectant dose. Our results showed that the ozone sensitivity distributions had higher uncertainties (**Figure S3**), leading to a wider range of the estimated values in **Figure 2C** and **Figure 3B**, because species’ data were limited as compared to the other two disinfectants.

**Fig 3.**
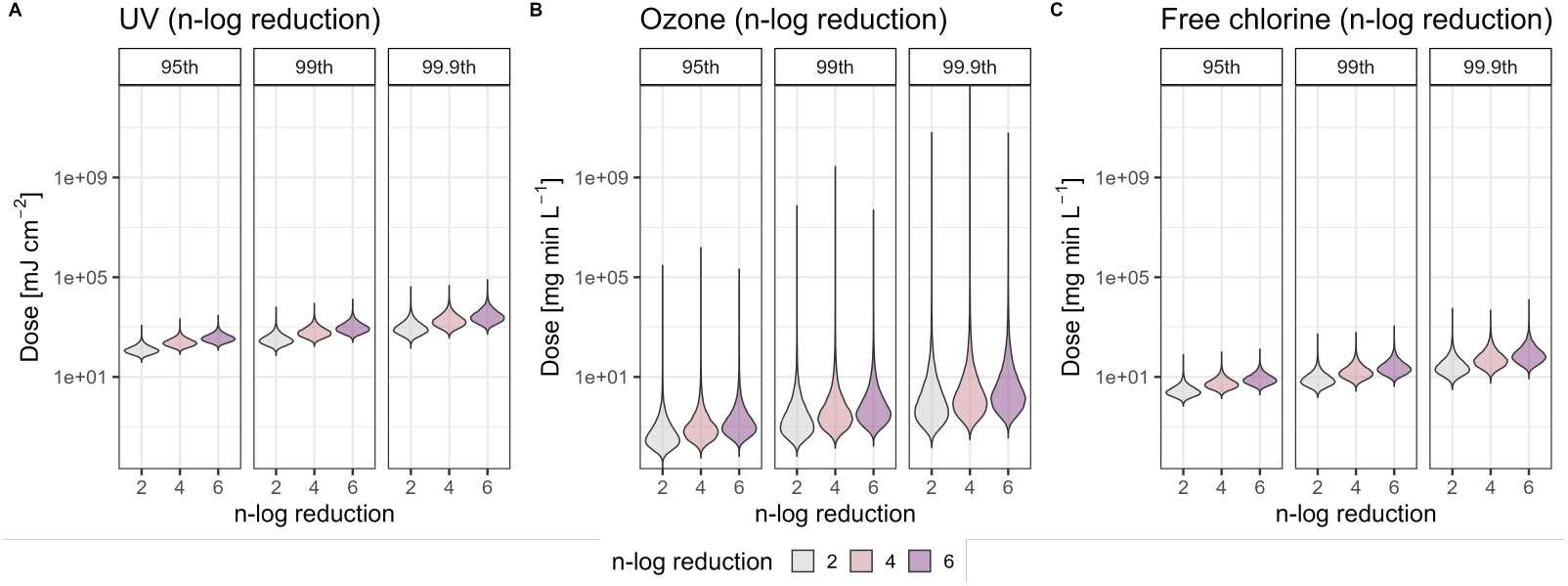
Required dose levels for UV (A), ozone (B), and free chlorine (C) to achieve each combination of the target log reduction value and the proportion of viruses expected to be inactivated. Light gray, light red, and purple plots represent 2-, 4-, 6-log reduction, respectively. Uncertainty in each estimate is presented as posterior distributions for each disinfectant, and the summary statistics of each posterior distribution is provided in Table S2 of Appendix.

## Discussion

In this study, we proposed a statistical framework to quantify the variation in sensitivity to standard disinfectants across different virus species by summarizing available experimental data on inactivation kinetics. This framework estimates disinfectant sensitivity distributions at the assemblage level, and translates a single reference value (e.g., log reduction values recommended in guidelines, or benchmark values referring to more resistant viruses found in recent experiments) into a proportion of virus species expected to be inactivated.

Our proposed framework has several aspects that can complement a single reference approach. As this framework captures the variation in sensitivity of viruses as a distribution, it allows for quantitative evaluation of the proportion of virus species that do not achieve the target log-reduction. Recent modelling studies in environmental microbiology have mainly aimed to predict inactivation rate constants under particular experimental conditions, and have elucidated individual-level (single virus- and condition-specific) inactivation kinetics for various viruses.^34^ By contrast, our approach summarizes such individual estimates as an overall disinfectant sensitivity distribution at the assemblage level. A contribution of the present study is to provide a reasonable way of extrapolating the data of tested viruses into those untested - when unidentified viruses are to be tested in the future, as long as the viruses are deemed to be randomly identified from the range of species currently covered, such viruses would be likely to follow the same disinfectant sensitivity distribution. Another strength of our Bayesian approach over a common inference procedure with maximum likelihood estimation is its capability of capturing the uncertainty in observed data. The present study treated observations from multiple experiments as interval data, rather than assuming that there is a set of single correct observations, and incorporated the variation in observed inactivation rate constants across different studies. Our framework provides a seamless and natural way of synthesizing available experimental data, and importantly, quantifying the uncertainty in required dose levels for an arbitrarily selected proportion of virus species to be inactivated as a posterior distribution (e.g., for a virus at the 95th percentile, shown in **Figure 3**).

Our motivation for this analysis is to make the extrapolation to untested viruses more reasonable, using the best available data. We inferred the distributions based on the species that have been tested for virological or operational reasons in the past,^3,23,24^ expecting currently untested viruses to be tested for the same or equivalent reason in the future. Using the exhaustive data contributes to more robust extrapolation across species by covering various virus species – our dataset also includes non-pathogenic viruses, which may enhance the extrapolation by borrowing the observed inactivation kinetics from viruses with similar virological characteristics, given that they have been tested as a surrogate of pathogenic viruses. The same arguments on the selection of species to construct the sensitivity distributions have been addressed in ecotoxicology, and many studies have proposed methodologic approaches and data-driven criteria,^19,22,35^ which are now used to form the current guidelines for ecological risk assessment. For example, the number of taxonomic groups (e.g., algae, invertebrates, fish, and amphibians) is often used to guarantee sufficient coverage of the species from different trophic levels. Such existing methodologies or discussions in ecotoxicology may help future research in environmental virology to identify important factors when estimating disinfectant sensitivity distributions. Our study has several limitations and suggestions for future work. The estimation of disinfectant sensitivity distributions requires observed data covering a sufficient number of species (five or more recommended as the minimum number of species in ecotoxicology).^22^ Although our present analysis covered a relatively large number of virus species, further accumulation of disinfection experiment data, in particular for ozone, would enhance the robustness of the estimation of distributions. Second, while our analysis focused only on a species-level classification, further stratification into subspecies-level or even grouping into a higher taxonomic level (e.g., genus-level) is possible. The granularity of this classification should be determined by the size or characteristic of the assemblage that the disinfectant sensitivity distribution aims to represent. For example, further stratification into a subgroup of viruses with capsid protein and the other may show a difference in the disinfectant sensitivity distributions for free chlorine. Third, our approach does not perfectly incorporate all the uncertainty and bias in the pooled inactivation rate constants. For instance, if there is a tailing effect in the relationship between disinfectant dose and log inactivation, that may lead to a biased estimate of inactivation constant. While our Bayesian approach partially incorporated the uncertainty in estimates between experiments, the bias due to the tailing effect should be further examined as the degree of tailing may differ by the type of virus and disinfectant.^36^

In conclusion, our proposed framework reveals the variation in sensitivity to UV, ozone, and free chlorine across different types of viruses, and quantifies the proportion of viruses expected to be inactivated given a disinfection condition of interest. This approach is simple and interpretable, which helps clarify the rationale of determining the required level of disinfectant doses - for instance, if our goal is set to inactivate 95% of species, we can obtain the required dose level based on estimated disinfectant sensitivity distributions, and such reasoning is easier to communicate with stakeholders. The use of disinfectant sensitivity distributions would complement, and provide supportive tools for probabilistic risk assessments and more transparent risk communication.

## Acknowledgements

The authors thank Dr. Leo Posthuma (The Dutch National Institute for Public Health and the Environment; Radboud University) for his invaluable input in improving the manuscript.

## Data and materials availability

All codes and simulated data are available at the authors’ GitHub link (https://github.com/miinay/disinfection_sensitivity_distribution).

