## Appendix for "Statistical framework for assessing heterogeneous sensitivity of viruses to ultraviolet, ozone, and free chlorine"

**Supplementary Materials**

**Supplementary text**

Data collection

Inactivation rate constant and log reduction value

Transforming guideline values

Bayesian framework to estimate parameter sets of distributions

**Other Supplementary Materials**

**Figure S1.** Histograms of reported inactivation rate constant *k* transformed according to 4 log reduction

**Figure S2.** Comparison of three fitted sensitivity distributions for UV

**Figure S3.** Comparison of three fitted sensitivity distributions for ozone

**Figure S4.** Comparison of three fitted sensitivity distributions for free chlorine

**Table S1.** Summary statistics of the estimated three parametric distributions and computed goodness-of-fit for 4-log reduction **Table S2.** Estimated dose levels required to achieve the inactivation of 95, 99, and 99.9 percent of viruses

**Supplementary text**

**Data collection**

The 220, 31, and 82 data on inactivation rate constants are collected from systematic reviews pertaining to UV,^1^ ozone,^2^ and free chlorine,^3^ respectively. The rate constants for UV were determined from a linear portion of the inactivation curve and did not include regions where tailing occurred.^1,3^ They were not screened by the conditions of pH or temperature due to their minimal effects on inactivation.^4^ Those for ozone were experimentally determined under a pH of 6.5–8.5 and a temperature of 15–25℃. Those for free chlorine are the harmonized values under a specific condition, a pH of 7.53, a temperature of 20 °C, and a chloride concentration of < 50 mM, taking the inter-study differences in physico-chemical conditions into account. The rate constants used for harmonization were estimated from the linear portion of the inactivation, excluding the tailing part. The processed dataset consists of 37, 9, and 28 species for UV, ozone, and free chlorine disinfection experiments, and is used for further analysis. While the majority of included studies reported mean and standard deviation of the estimated inactivation rate constants, several studies reported only mean values. To enhance the statistical power in determining sensitivity distributions, we also included those mean values and treated them as a point estimate without any uncertainty interval. The summary of analyzed data with details (i.e., inoculation methods and references) is provided in the author's GitHub repository (https://github.com/miinay/disinfection_sensitivity_distribution).

**Inactivation rate constant and log reduction value**

To synthesize the obtained data on inactivation rate constants, we set two assumptions. First, the reported inactivation rate constant $k$ was converted into an arbitrary log reduction value (with the reduction order of $n$) by assuming the pseudo-first-order decay relationship (i.e., consistent with the Chick-Watson model, where the dilution coefficient is set to be one^5,6^). The survival proportion of disinfected virus $S_{dis}(d)$ given the disinfectant dose $d$ is expressed as:

$$S_{dis}\left( d \right)=exp \left( -kd \right). \#Eq.S1$$

This equation allows for transforming an estimate of an inactivation rate constant $k$ into an arbitrary log reduction value $n$. The relation is, by definition,

$$S_{dis}\left( d_{{LRV}_{n}} \right)={10}^{-n}=exp \left( -kd_{{LRV}_{n}} \right). \#Eq.S2$$

and this reduces to

$$d_{{LRV}_{n}}=\frac{n}{k}ln10 \#Eq.S3$$

where $d_{{LRV}_{n}}$ is the corresponding required dose of disinfectants to achieve $n$ log reduction.

Second, to incorporate the uncertainty in the range of reported $k$, we took the lower and upper values of 95% confidence intervals $\left[ k_{lower}, k_{upper} \right]$, and treated them as interval data, assuming that a value $k$ is uniformly distributed within this interval. This interval was converted into the unit of required dose of disinfectants, using **Eq. S3**.

**Transforming guideline values**

Disinfectant doses required for a certain level of inactivation were determined by existing USEPA Guidance manuals for the Long Term 2 Enhanced Surface Water Treatment Rule.^7^

*UV*

Given that UV dose of 186 mJ cm^-2^ is recommended to achieve viral inactivation at 4 log, required dose for n log inactivation was generalized to be 46.5 × n (mJ cm^-2^).^7^

*Ozone*

The USEPA CT model for ozone disinfection of viruses at n-log is given by 2.1744 × 1.0726^Temperature^ × CT.^2^ Required CT for n-log inactivation at 20 °C is thus 0.113 × n (mg min L^-1^).

*Free chlorine*

Given that CT values of 3 mg min L^-1^ are recommended to achieve viral inactivation at 4 log at pH 6–9 and at 20 °C, required CT values for n-log inactivation is set to be 0.75 × n (mg min L^-1^).^8^

**Bayesian framework to estimate parameter sets of distributions**

We fitted three parametric distributions to the observed inactivation rate constants (after converting them into the corresponding disinfection dose), using a likelihood-based approach that allows for reported value range to be a single point estimate or an interval.^9,10^ Three parametric distributions, which have been widely used in ecotoxicology and ecological risk assessment, were examined: lognormal, Weibull, and gamma distributions.

Our likelihood-based approach uses a Bayesian framework to estimate the set of parameters for each distribution $\theta$.^9,10^ We specified positive flat prior distributions, used the default of 4 chains each with 3000 burn-in and 10,000 sampling steps. For all models, convergence was assessed using the Rhat diagnostic.^11^ We computed the posterior distributions of each parameter using the {rstan} package in the R statistical programming environment version 4.1.3.^12^

**Supplementary figures and tables**

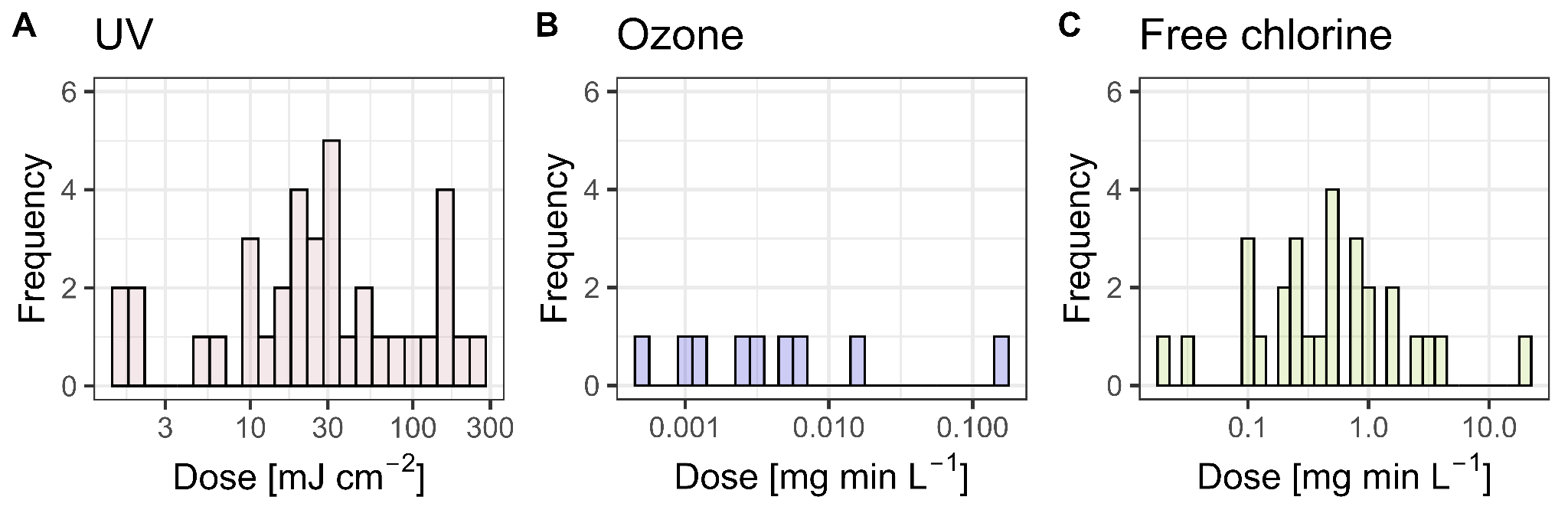

**Figure S1.** Histograms of the transformed mean inactivation rate constants where 4-log reduction is set as a goal

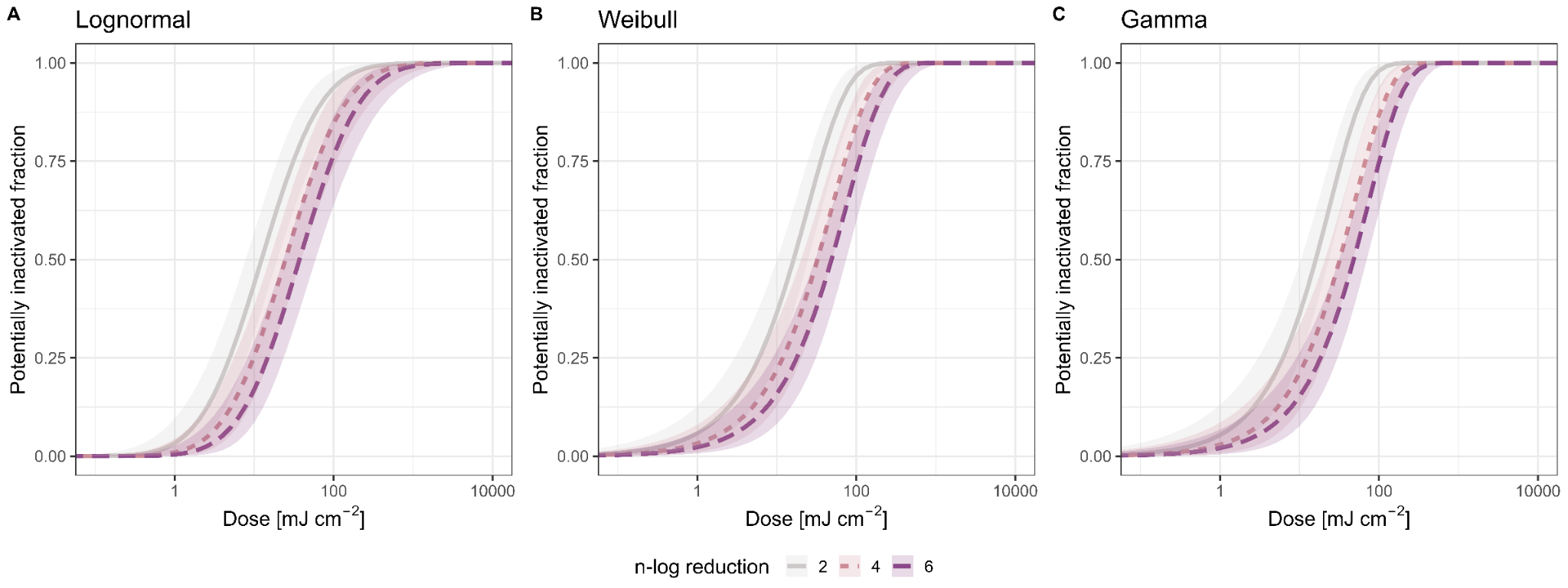
**Figure S2.** Estimated disinfectant sensitivity distributions for UV using lognormal (A), Weibull (B), and gamma distributions (C). Light gray, light red, and purple curves represent 2-, 4-, 6-log reduction, respectively. The solid lines are the estimated disinfectant sensitivity distributions, and the shaded areas indicate 95% credible intervals computed using posterior samples.

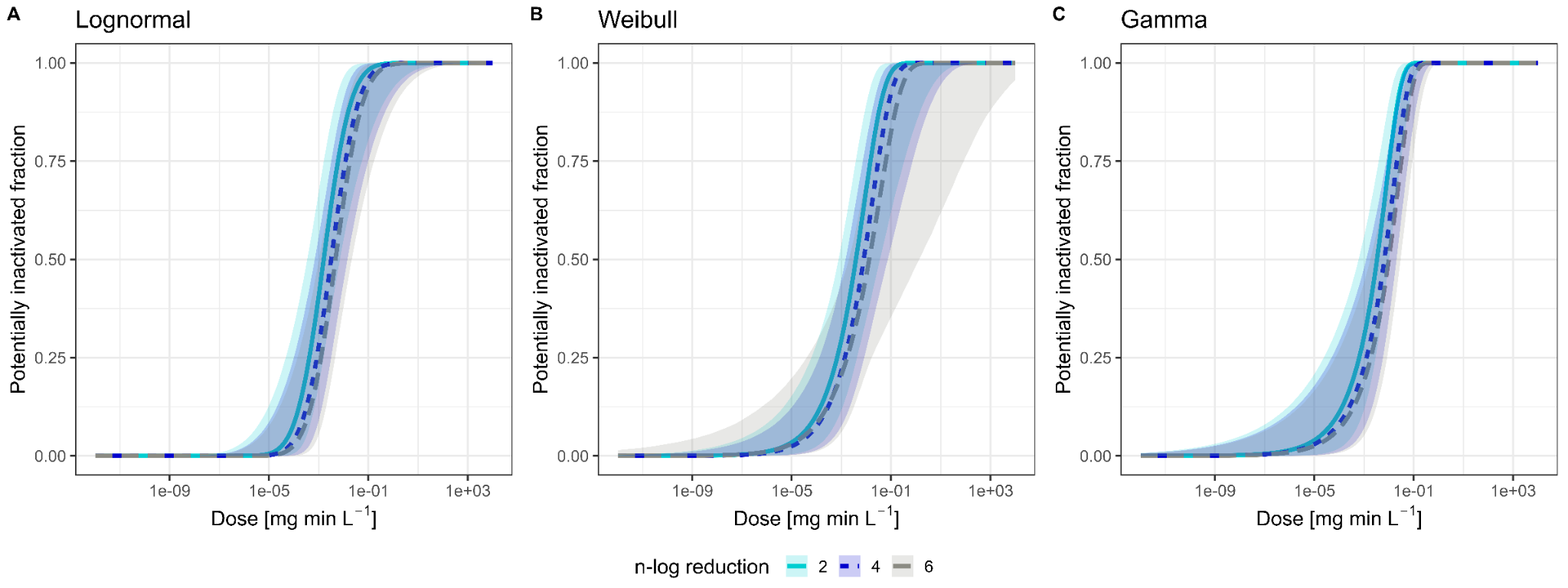

**Figure S3.** Estimated disinfectant sensitivity distributions for ozone using lognormal (A), Weibull (B), and gamma distributions (C). Light blue, darker blue, and gray curves represent 2-, 4-, 6-log reduction, respectively. The solid lines are the estimated disinfectant sensitivity distributions, and the shaded areas indicate 95% credible intervals computed using posterior samples.

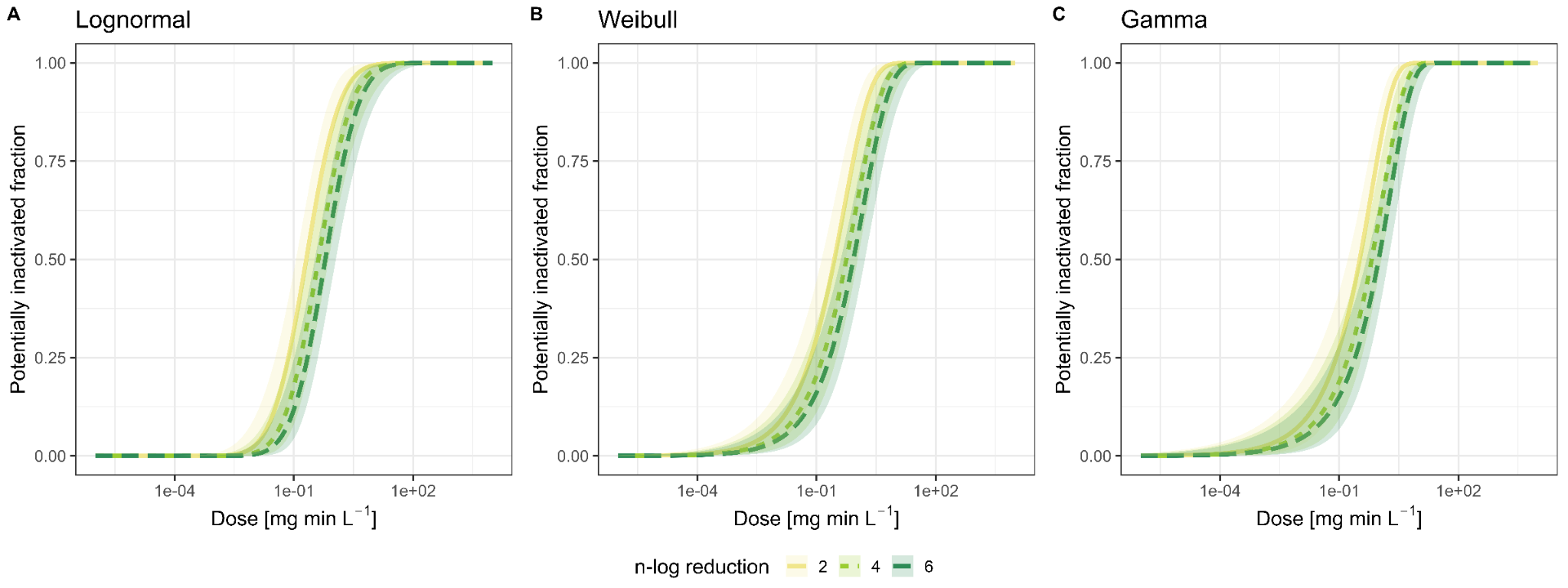

**Figure S4.** Estimated disinfectant sensitivity distributions for free chlorine using lognormal (A), Weibull (B), and gamma distributions (C). Yellow, light green, and dark green curves represent 2-, 4-, 6-log reduction, respectively. The solid lines are the estimated disinfectant sensitivity distributions, and the shaded areas indicate 95% credible intervals computed using posterior samples.

**Table S1.** Summary statistics of the estimated three parametric distributions and computed goodness-of-fit for 4-log reduction

|  |  | Mean | SD | WAIC^*^ | LOOIC^†^ |
| --- | --- | --- | --- | --- | --- |
| UV | Lognormal | 63.3 [95%CrI^‡^: 37.0–141.9] | 147.8 [95%CrI: 61.2–624.8] | 370.8 | 371.7 |
|  | Gamma | 48.6 [95%CrI: 34.9–71.4] | 51.2 [95%CrI: 35.6–81.3] | 369.8 | 371.3 |
|  | Weibull | 52.9 [95%CrI: 36.7–82.4] | 60.0 [95%CrI: 39.4–112.7] | 369.6 | 370.8 |
| Ozone | Lognormal | 0.024 [95%CrI: 0.0044–8.06] | 0.17 [95%CrI: 0.011–1.37×10^4^] | -62.5 | -61.5 |
|  | Gamma | 0.0162 [95%CrI: 0.0069–0.0498] | 0.0241 [95%CrI: 0.011–0.0835] | -53.9 | -51.6 |
|  | Weibull | 0.031 [95%CrI: 0.0079–0.971] | 0.0642 [95%CrI: 0.0145–5.62] | -58.3 | -56.7 |
| Free chlorine | Lognormal | 1.35 [95%CrI: 0.66–4.58] | 4.18 [95%CrI: 1.28–36.8] | 58.2 | 59.3 |
|  | Gamma | 1.31 [95%CrI: 0.82–2.27] | 1.74 [95%CrI: 1.09–3.25] | 74.0 | 76.1 |
|  | Weibull | 1.39 [95%CrI: 0.79–2.90] | 2.21 [95%CrI: 1.19–5.93] | 66.4 | 68.2 |

^*^WAIC: Widely applicable information criterion. ^†^LOOIC: Leave-One-Out Information Criterion; these values indicate the goodness-of-fit, where lower values indicate a better fit. ^‡^CrI, credible interval.

**Table S2.** Estimated dose levels required to achieve the inactivation of 95, 99, and 99.9 percent of viruses.

|  |  | 95th percentile | 99th percentile | 99.9th percentile |
| --- | --- | --- | --- | --- |
| UV | 2-log | 1.17×10^2^ [95%CrI: 63.2–2.75×10^2^] | 2.96×10^2^ [95%CrI: (1.38–8.91) ×10^2^] | 8.40×10^2^ [95%CrI: 3.26×10^2^–3.41×10^3^] |
|  | 4-log | 2.34×10^2^ [95%CrI: (1.28–5.42) ×10^2^] | 5.94×10^2^ [95%CrI: 2.79×10^2^–1.76×10^3^] | 1.70×10^3^ [95%CrI: 6.52×10^2^ –6.66×10^3^] |
|  | 6-log | 3.51×10^2^ [95%CrI: (1.92–8.12×10^2^] | 8.92×10^2^ [95%CrI: 4.17×10^2^ –2.64×10^3^] | 2.54×10^3^ [95%CrI: 9.78×10^2^ –1.00×10^4^] |
| Ozone | 2-log | 0.04 [95%CrI: 0.01–1.22] | 0.16 [95%CrI: 0.02 –13.7] | 0.75 [95%CrI: 0.06– 240] |
|  | 4-log | 0.08 [95%CrI: 0.02–3.02] | 0.33 [95%CrI: 0.04 –38.8] | 1.5 [95%CrI: 0.12–702] |
|  | 6-log | 0.12 [95%CrI: 0.02–4.33] | 0.49 [95%CrI: 0.06 –52.3] | 2.25 [95%CrI: 0.18– 933] |
| Free chlorine | 2-log | 2.55 [95%CrI: 1.19– 8.14] | 7.29 [95%CrI: 2.81 –32.1] | 23.6 [95%CrI: 7.17 –154] |
|  | 4-log | 5.12 [95%CrI: 2.37–16.2] | 14.6 [95%CrI: 5.58–64.3] | 47.5 [95%CrI: 14.3 –304] |
|  | 6-log | 7.7 [95%CrI: 3.59–23.9] | 22.0 [95%CrI: 8.42–94.5] | 71.3 [95%CrI: 21.6–448] |
